# Spatial memory performance is associated with region-specific coordination of hippocampo–cortical sleep oscillations

**DOI:** 10.64898/2026.01.27.702052

**Authors:** Mauricio Caneo, Nelson Espinosa, Alejandro Aguilera, Guillermo Lazcano, Ariel Lara-Vasquez, Pablo Fuentealba

## Abstract

Sleep plays a critical role in memory consolidation, yet how coordination among sleep oscillations across hippocampo–cortical circuits relates to behavioral performance remains incompletely understood. Here, we examined how cross-regional coordination of cardinal sleep oscillations during non–rapid eye movement (nREM) sleep relates to spatial memory performance. Adult rats were trained in an object–place recognition task and allowed to sleep while local field potentials were recorded simultaneously from dorsal and ventral hippocampus (CA1d and CA1v) and retrosplenial, prefrontal, and lateral entorhinal cortices. We first confirm that nREM sleep duration, but not REM sleep, is positively associated with spatial memory performance. We then characterized the temporal relationships between hippocampal sharp-wave ripples and cortical spindles and slow oscillations, revealing region- and oscillation-specific coordination patterns that depend on the dorso–ventral origin of hippocampal ripple events. Finally, by comparing task-related changes in ripple-triggered cortical activity between low- and high-performing sessions, we identify selective, performance-dependent modulation of hippocampo–cortical coupling during task-related sleep. Spatial memory performance was consistently associated with enhanced ripple–spindle and ripple–slow oscillation coupling driven by dorsal hippocampal ripples, particularly in interactions with entorhinal and prefrontal cortices, whereas ventral hippocampal interactions failed to show behavioural association. Together, these results reveal that spatial memory performance is linked to region- and timing-specific coordination of sleep oscillations across hippocampo–cortical networks during nREM sleep, highlighting a functional differentiation of dorsal and ventral hippocampal contributions to sleep-dependent memory consolidation.

## Introduction

Sleep is a fundamental biological process that supports the stabilization and reorganization of newly acquired memories [1,2]. Converging evidence from human and animal studies indicates that memory consolidation is particularly dependent on non–rapid eye movement (nREM) sleep [3,4], a brain state characterized by the coordinated expression of distinct neuronal oscillations across distributed brain networks [5]. Among these, cortical slow oscillations [6], thalamocortical sleep spindles [7], and hippocampal sharp-wave ripples [8] are considered cardinal sleep rhythms and are widely implicated in systems-level memory processing. While each of these oscillations has been independently linked to memory consolidation, growing evidence suggests that their precise temporal coordination across hippocampo–cortical circuits is critical for effective memory storage [9–11].

The hippocampus plays a central role in spatial memory [8,12] and interacts extensively with neocortical regions during both learning and sleep [9,13,14]. During nREM sleep, hippocampal sharp-wave ripples are thought to support memory consolidation by reactivating recently encoded information and promoting its transfer to cortical networks [15–17]. This process is proposed to be facilitated by the temporal alignment of ripples with cortical slow oscillations and sleep spindles, which provide windows of heightened synaptic plasticity [18]. Disruption of ripple–spindle or ripple–slow oscillation coupling impairs memory consolidation, underscoring the importance of cross-regional oscillatory coordination [11,15]. However, how such coordination is organized across different cortical targets, and how they relate to behavioral performance, remains incompletely understood.

An additional level of complexity arises from the functional heterogeneity along the dorso-ventral axis of the hippocampus [19,20]. The dorsal hippocampus is preferentially involved in spatial and cognitive processing [21,22], whereas the ventral hippocampus is more strongly associated with affective and motivational functions [19,23,24]. Anatomical and functional studies indicate that dorsal and ventral hippocampal subregions exhibit distinct connectivity patterns with cortical areas [20], suggesting that sleep-dependent hippocampo–cortical coordination may differ depending on the hippocampal source of ripple activity [25]. Despite this, most studies have treated hippocampal ripples as a unitary phenomenon, without explicitly considering longitudinal specialization.

Furthermore, although several studies have demonstrated correlations between sleep oscillations and memory performance [26–28], fewer have directly linked individual differences in behavioral outcome to task-related changes in cross-regional sleep connectivity. Understanding how performance-dependent modulation of hippocampo–cortical coordination emerges during post-task sleep is essential for elucidating the network mechanisms that support successful memory consolidation.

In the present study, we addressed these gaps by examining how cross-regional coordination of sleep oscillations during nREM sleep relates to spatial memory performance. Using a spatial object–place recognition task [29–31] in rats, combined with simultaneous recordings from dorsal and ventral hippocampus and multiple interconnected cortical regions, we first assessed the relationship between sleep architecture and behavioral performance. We then characterized the temporal dynamics of ripple-triggered cortical spindles and slow oscillations, explicitly comparing coupling driven by dorsal versus ventral hippocampal ripples. Finally, we asked whether task-related changes in hippocampo–cortical coupling during sleep differentiate low- and high-performing animals. By integrating behavioral, anatomical, and electrophysiological analyses, this work provides a network-level perspective on how distributed sleep oscillations support spatial memory consolidation.

## Results

### Non-REM sleep duration is positively associated with spatial memory performance

We trained adult rats in the object–place recognition (OPR) task, a well-established assay of spatial memory [29,32] based on object exploration (Table S1), and allowed animals to sleep ad libitum during two fixed resting periods; one preceding the behavioral task and one occurring during between task phases (Table S2, **Fig. 1A**). Animals were chronically implanted with electrodes targeting the dorsal and ventral hippocampus (CA1d and CA1v, respectively) as well as associated cortical regions, comprising the retrosplenial cortex (RSC), prefrontal cortex (PFC), and lateral entorhinal cortex (LEC) (Table S3, Fig. S1). Neural activity was continuously recorded across the sleep–wake cycle and throughout task performance. Consistent with previous work, we observed that the cumulative duration of spontaneous sleep strongly associated with spatial discrimination during the retrieval phase, as longer total sleep intervals were positively correlated with higher preference indices, indicative of superior spatial memory recall (Fig. S2). To characterize vigilance states, we combined local field potential recordings with video-based monitoring of body movements [3,33], allowing us to construct hypnograms for each resting session and to reliably distinguish sleep stages (**Fig. 1B**). Rapid eye movement (REM) sleep was identified by sustained hippocampal theta oscillations during prolonged periods of behavioral immobility (**Fig. 1C**), whereas non-REM (nREM) sleep was characterized by prominent hippocampal ripple events in conjunction with cortical spindle oscillations (10–16 Hz; **Fig. 1D**) or slow-wave activity (0.5–4 Hz; Fig. S3). A linear mixed-effects model, with brain state included as a fixed effect and animal identity and recording session as random effects, revealed that time spent in REM sleep was not associated with spatial memory performance (P = 0.535; **Fig. 1E**). In contrast, nREM sleep duration showed a significant positive association with retrieval accuracy (P = 0.033; **Fig. 1F**). Consistent with these results, only nREM sleep emerged as a significant predictor of memory performance in the model (Table S4). Together, these results are consistent with prior observations in both humans and rodents, reinforcing the specific contribution of nREM sleep to spatial memory consolidation [3,34,35].

**Figure 1.**
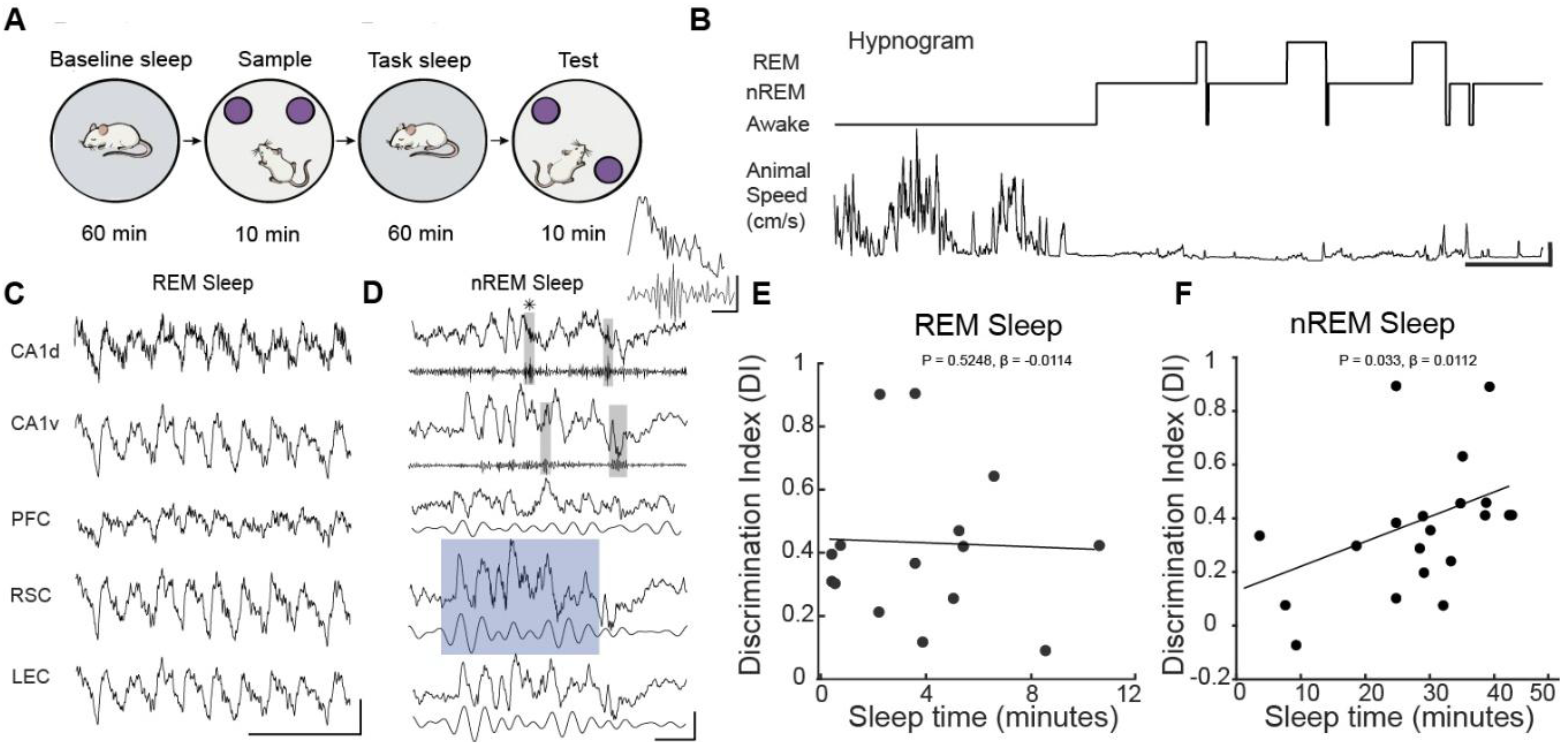
Relationship between sleep architecture and spatial memory performance. A, Experimental timeline illustrating baseline sleep, object–place recognition (OPR) task, and task-related sleep. Animals slept ad libitum on a retention platform before and after the behavioral task. B, Example hypnogram (rat AA04, session 1) showing brain state classification into quiet wakefulness (QW), non–rapid eye movement (nREM) sleep, and rapid eye movement (REM) sleep. Scale bars: horizontal, 500 s; vertical, 5 cm/s. C, Example simultaneous local field potential (LFP) recordings from hippocampus (CA1d, CA1v) and cortex (PFC, RSC, LEC) illustrating widespread, sustained theta oscillations used to identify REM sleep (rat AA07, session 1). Scale bars: horizontal, 0.5 s; vertical, 1 mV. D, Example LFP traces showing hippocampal sharp-wave ripples (gray shading) and concomitant cortical spindle activity (blue shading) during nREM sleep (rat AA07, session 1). Scale bars: horizontal, 100 ms; vertical, 0.5 mV. Inset, example ripple expanded and depicted by asterisk (*). Scale bars: horizontal, 50 ms; vertical, 0.5 mV. E, Relationship between REM sleep duration and discrimination index, showing no significant association (LMM, P = 0.525, β = -0.01;). F, Positive association between nREM sleep duration and spatial memory performance, quantified by the discrimination index (LMM, P = 0.033, β = 0.01; n = 10 rats, 20 sessions). Linear mixed model (LMM), Discrimination Index ∼ REM duration + nREM duration + (1|animal).

### Cross-regional coordination of sleep oscillations in the hippocampo– cortical circuit

Cortical slow waves, thalamic spindles, and hippocampal sharp-wave ripples are considered cardinal sleep oscillations and are thought to play a central role in memory consolidation [36]. Accordingly, we examined the cross-regional dynamics of these oscillations during nREM sleep. To assess the temporal relationship between hippocampal ripples and cortical spindle activity, we quantified spindle probability in cortical regions aligned to ripple onset in CA1d (**Fig. 2A**). Cortical spindle probability progressively increased preceding dorsal hippocampal ripple occurrence, and was significantly different from zero for RSC and LEC, but not for PFC (Fig. S4). This pre-ripple buildup was larger in RSC (3.47 ± 0.18 z) than LEC (2.55 ± 0.25 z, P = 7×10^-7^; **Fig. 2B**). However, peak latencies did not differ across regions (P = 0.6; **Fig. 2C**). Immediately following ripple onset, spindle probability dropped sharply in both regions. We next repeated this analysis using ripple events detected in CA1v as the temporal reference, enabling direct comparison with CA1d-aligned spindle dynamics (**Fig. 2D**). Cortical spindle probability also increased preceding ventral hippocampal ripple occurrence, and was significantly different from zero for RSC and PFC, but not for LEC (Fig. S4). Spindle probability rose gradually with similar kinetics preceding the ripple onset, reaching comparable peak amplitudes between RSC and PFC (P = 0.18; **Fig. 2E**), but different peak latencies (P = 1.2×10^-6^; **Fig. 2F**). Quantification of these effects using linear mixed-effects models confirmed significant region-specific differences in spindle amplitude and latency (Table S5). Thus, CA1v-aligned spindle modulation showed reduced regional differentiation. Following ripple onset, spindle probability rapidly decreased in all regions.

**Figure 2.**
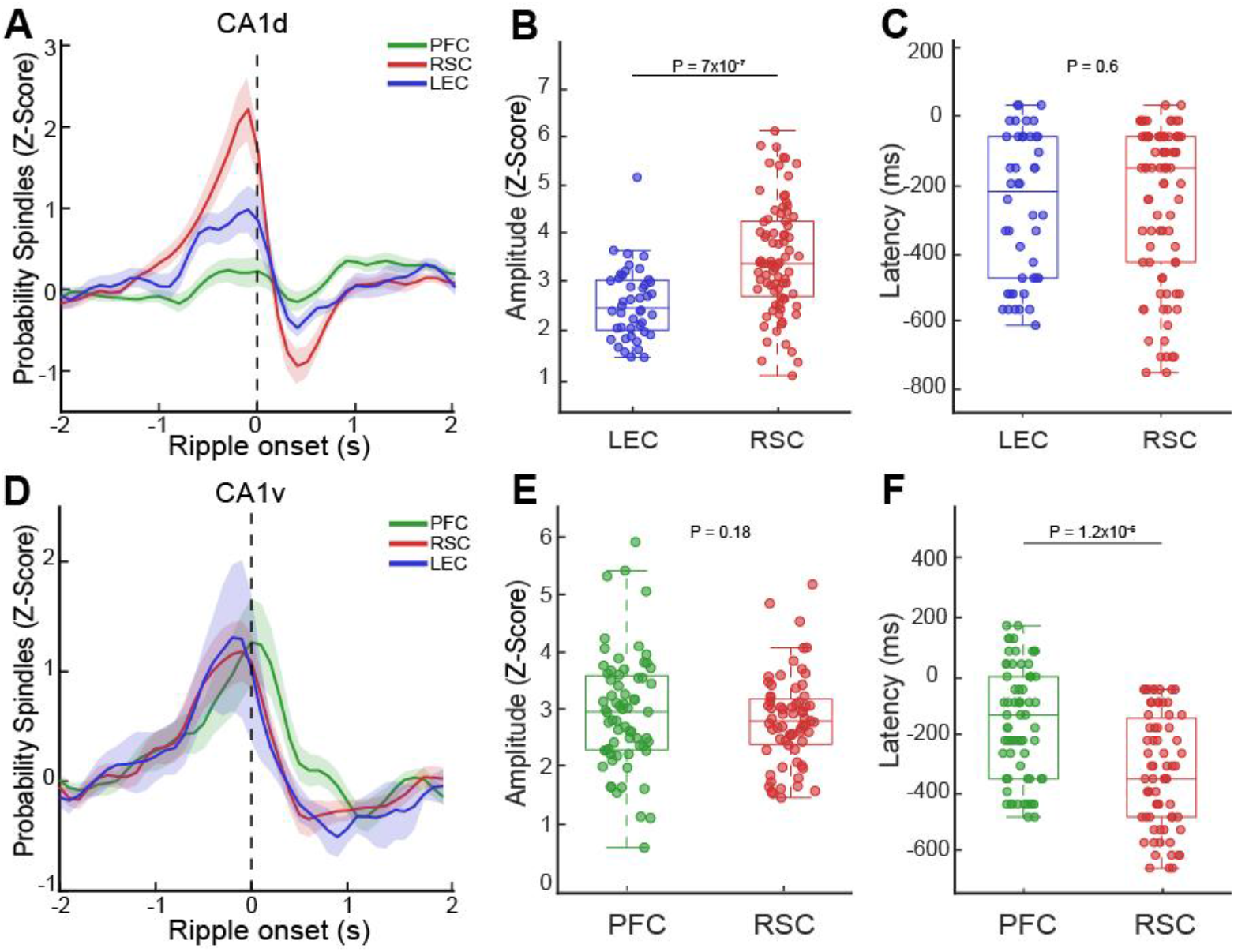
Regional coordination of cortical spindle activity with hippocampal ripples. A, Ripple-triggered cortical spindle probability aligned to ripple onset in CA1d for RSC, LEC, and PFC. B, Peak spindle probability preceding CA1d ripples across cortical regions (LMM, P = 7×10^-7^). C, Latency of peak spindle probability relative to CA1d ripple onset (LMM, P = 0.6). D, Ripple-triggered cortical spindle probability aligned to ripple onset in CA1v. E, Peak spindle probability preceding CA1v ripples(LMM, P = 0.18). F, Latency of peak spindle probability relative to CA1v ripple onset (LMM, P = 1.2×10^-6^). Shaded areas indicate ± SEM. Differences in amplitude and latency were assessed using linear mixed-effects models (LMM): Crosscorrelagram amplitude ∼ target areas + (1|animal), and Crosscorrelagram latency ∼ target areas + (1|animal).

We next examined the temporal relationship between hippocampal ripples and cortical slow oscillation (SO) probability, again using hippocampal ripples as the temporal reference. When aligned to CA1d ripples, SO probability exhibited marked regional differences around ripple onset (**Fig. 3A**). Cortical SO probability in relation to dorsal hippocampal ripples was significantly different from zero for RSC and PFC, but not for LEC (Fig. S4). In RSC, SO probability increased sharply near the ripple onset, reaching larger peak amplitude (3.33 ± 0.24 z) than PFC (2.34 ± 0.31 z, P = 6.1×10^-5^; **Fig. 3B**). Peak latencies were different, as RSC SO peaked before ripple onset (-289.99 ± 48.48 ms), while PFC SO occurred significantly later following ripple initiation (199.40 ± 12.61 ms, P = 8.15 ×10^-38^; **Fig. 3C**). When ripples were instead referenced to CA1v (**Fig. 3D**), SO peak probability was significantly different from zero in RSC and LEC, but not in PFC (Fig. S4). In this case, both peak amplitude (P = 6.2×10^-4^; **Fig. 3E**) and peak latency (P = 4.6×10^-6^; **Fig. 3F**) were larger in LEC when compared to RSC. Across both reference conditions, post-ripple SO probability gradually returned toward baseline. Linear mixed-effects modeling revealed significant target-region effects on ripple-triggered slow oscillation amplitude and timing (Table S6).

**Figure 3.**
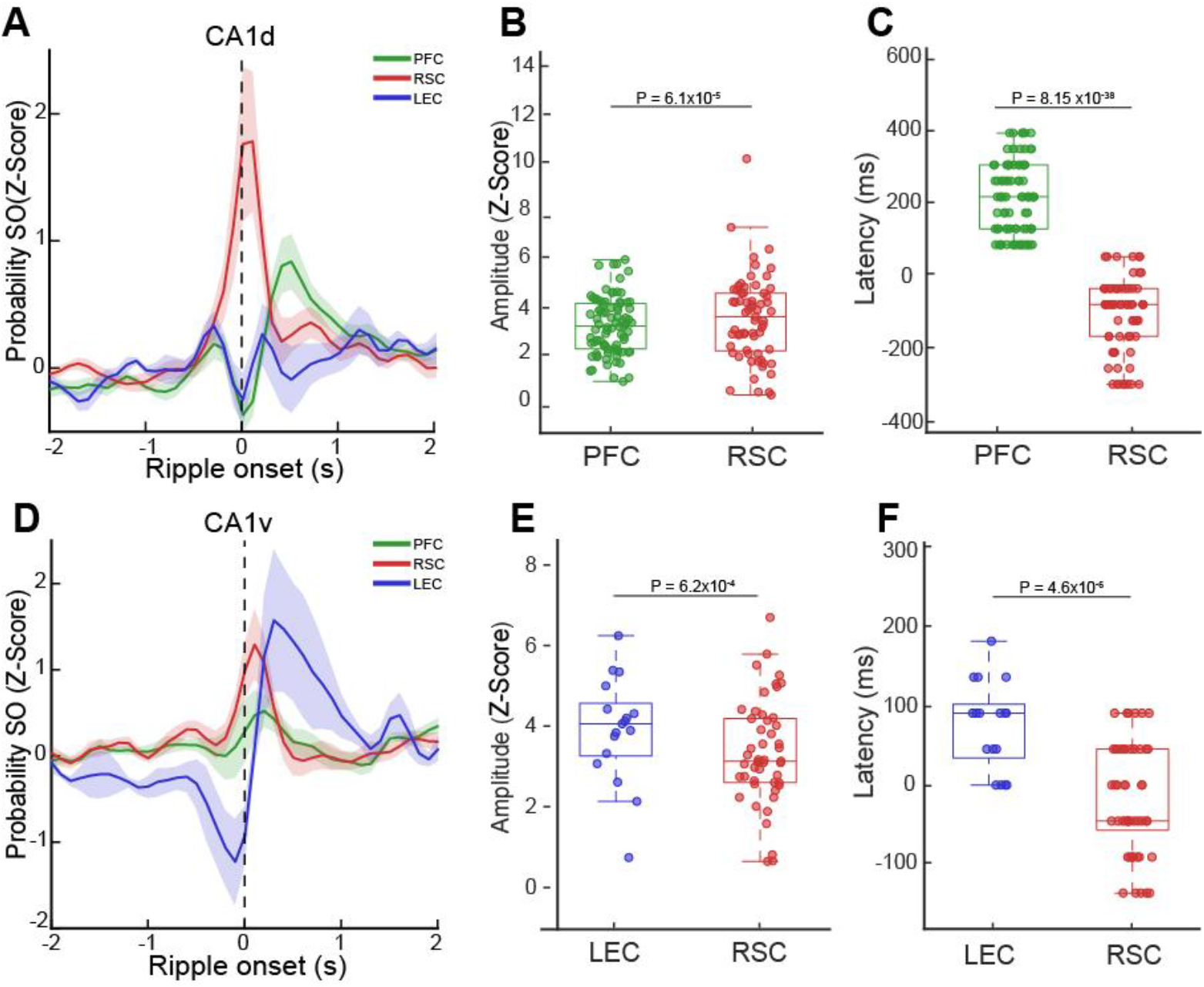
Regional coordination of cortical slow oscillations with dorsal and ventral hippocampal ripples. A, Ripple-triggered cortical slow oscillation (SO) probability aligned to CA1d ripple onset. B, Peak SO probability for cortical regions referenced to CA1d ripples (LMM, P = 6.1×10^-5^). C, Latency of SO peak relative to CA1d ripple onset (LMM, P = 8.15×10^-38^). D, Ripple-triggered SO probability aligned to CA1v ripple onset. E, Peak SO probability for cortical regions referenced to CA1v ripples (LMM, P = 6.2×10^-4^). F, Latency of SO peak relative to CA1v ripple onset (LMM, P = 4.6×10^-6^). Shaded areas indicate ± SEM. Regional differences in SO amplitude and latency were quantified using linear mixed-effects models (LMM): Crosscorrelagram amplitude ∼ target areas + (1|animal), and Crosscorrelagram latency ∼ target areas + (1|animal).

Together, these results evidence that the temporal coordination of cortical spindles and SO during nREM sleep depends on the septo-temporal origin of hippocampal ripple events, revealing region-specific patterns of hippocampo– cortical coupling that may shape information flow across the hippocampo-cortical network.

### Spatial memory performance correlates with changes in cross-regional sleep connectivity

To relate hippocampo–cortical connectivity during sleep to spatial memory performance, we examined task-related changes in cross-regional coupling of sleep oscillations as a function of behavioral outcome in the OPR task. Owing to the limited sample size, experimental sessions were stratified into low- and top-performing groups using a median split of the discrimination index. For each animal and session, we quantified task-induced modulation of sleep connectivity by computing the difference between ripple-triggered cross-correlograms obtained during task-related sleep and those measured during baseline sleep. This subtraction isolated learning-related changes in sleep coupling while controlling for baseline connectivity. Analyses were restricted to region pairs and temporal windows that were previously identified as significantly modulated relative to baseline (Fig. S4). The resulting difference measures were then grouped by behavioural performance and compared across groups.

Overall, task-related changes in ripple-associated cortical coupling differentiated performance groups, but in a region- and timing-specific manner. We first examined hippocampal ripple–cortical spindle coupling using ripples detected in CA1d as the temporal reference (**Fig. 4A**). CA1d–LEC coupling showed a significant enhancement in spindle probability in top-performing sessions relative to low-performing sessions (P = 0.039; **Fig. 4B**). In contrast, CA1d– RSC spindle coupling did not differ between performance groups (P = 0.872; **Fig. 4C**), indicating selective engagement of the entorhinal pathway in successful memory performance. We next repeated the analysis using ripples detected in CA1v as the temporal reference (**Fig. 4D**). In this case, no performance-dependent differences were observed across cortical regions. Neither CA1v–PFC coupling (P = 0.356; **Fig. 4E**) nor CA1v–RSC coupling (P = 0.213; **Fig. 4F**) differed between performance groups, suggesting a limited contribution of ventral hippocampal ripples to performance-related spindle coordination. Performance-dependent effects were formally assessed using linear mixed-effects models comparing low- and top-performing sessions (Table S7). We then applied the same performance-based comparison to hippocampal ripple–cortical SO coupling (**Fig. 5A**). Using CA1d ripples as the temporal reference, we observed enhancement of task-related coupling in top-performing sessions for both CA1d–RSC (P = 0.0002; **Fig. 5B**) and CA1d–PFC (P = 0.028; **Fig. 5C**) connections. In contrast, when CA1v ripples were used as reference (**Fig. 5D**), no significant performance-related differences were detected. Indeed, neither CA1v–RSC (P = 0.076; **Fig. 5E**) nor CA1v–LEC coupling (P = 0.322; **Fig. 5F**) differed between groups. These performance-related differences in ripple– slow oscillation coupling were confirmed by linear mixed-effects modeling (Table S8).

**Figure 4.**
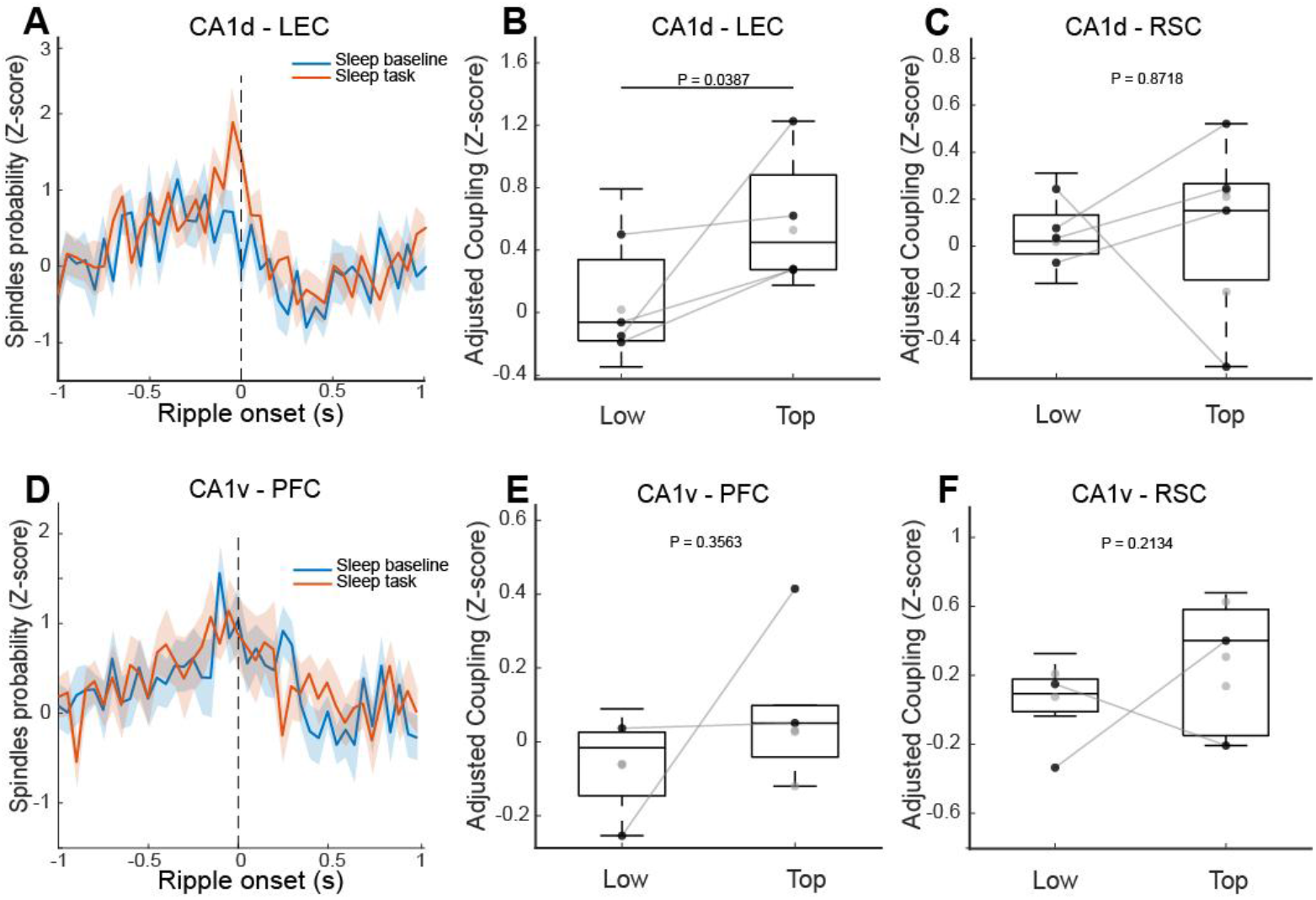
Performance-dependent modulation of ripple–spindle coupling during sleep. A, CA1d ripple-triggered LEC spindle probability illustrating task-related changes in ripple–spindle coupling, computed as the difference between task-related (red) and baseline (blue) sleep. B, CA1d–RSC spindle coupling does not differ between low- and top-performing sessions (LMM, P = 0.8718). C, Enhanced CA1d–LEC spindle coupling in top-performing sessions (LMM, P = 0.0387). D, CA1v ripple-triggered PFC spindle probability illustrating task-related changes in ripple–spindle coupling. E, CA1v–RSC spindle coupling shows no performance-dependent modulation (LMM, P = 0.2134). F, CA1v–PFC spindle coupling shows no performance-dependent modulation (LMM, P = 0.3563). Group comparisons were assessed using linear mixed-effects models (LMM): Delta sleep (task – baseline) ∼ group + (1|animal). Black circles depict sessions from the same animal (connected by gray lines), grey circles point to sessions from different animals.

**Figure 5.**
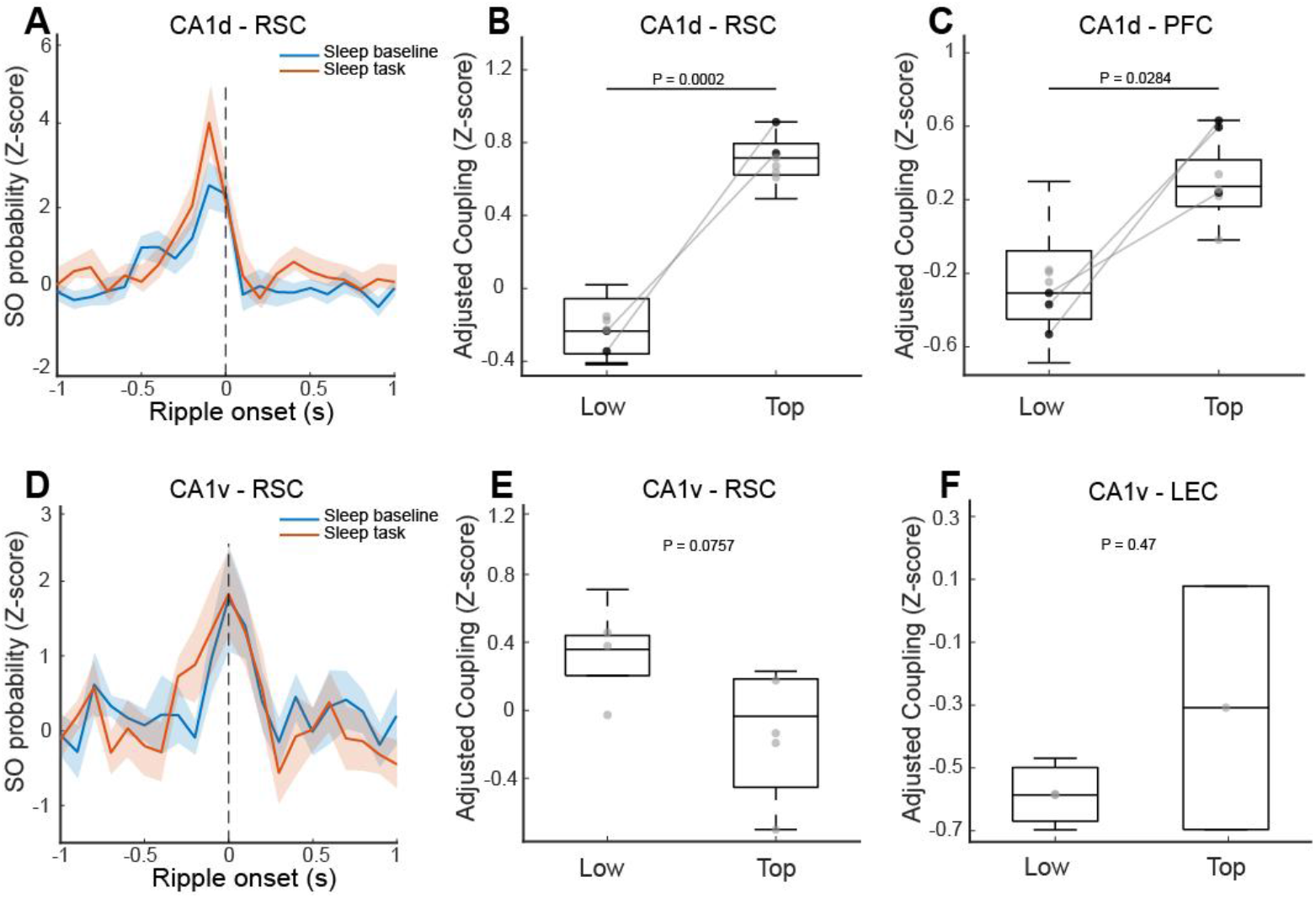
Performance-dependent modulation of ripple–slow oscillation coupling during sleep. A, CA1d ripple-triggered RSC slow oscillation probability illustrating task-related changes in ripple–slow oscillation coupling, computed as the difference between task-related (red) and baseline (blue) sleep. B, Enhanced CA1d–PFC ripple–SO coupling in top-performing sessions (LMM, P = 0.0284). C, Enhanced CA1d–RSC ripple–SO coupling in top-performing sessions (LMM, P = 0.0002). D, CA1v ripple-triggered RSC slow oscillation probability illustrating task-related changes in ripple–slow oscillation coupling. E, CA1v–RSC ripple–SO coupling does not differ between performance groups (LMM, P = 0.0757). F, CA1v–PFC ripple–SO coupling does not differ between performance groups (LMM, P = 0.3217). Statistical comparisons were performed using linear mixed-effects models (LMM) on task-related changes relative to baseline sleep: Delta sleep (task – baseline) ∼ group + (1|animal). Black circles depict sessions from the same animal (connected by gray lines), grey circles point to sessions from different animals.

Taken together, these results indicate that spatial memory performance is selectively associated with task-related, region- and timing-specific modulation of hippocampal ripple–cortical coupling during nREM sleep. Interestingly, successful memory performance is linked most consistently to connectivity supported by CA1d ripples, underscoring a privileged role for dorsal hippocampal output in coordinating sleep-dependent cortical processes that support spatial memory consolidation.

## Discussion

Our study provides new insight into the systems-level mechanisms by which hippocampo–cortical coordination during sleep contributes to spatial memory consolidation. By tracking regional coordination across the hippocampal longitudinal axis (CA1d and CA1v) and multiple cortical targets (RSC, PFC, LEC), we identify dissociable routes of information flow and uncover the behavioral relevance of specific ripple–spindle–SO coupling motifs. These findings support and refine contemporary models of memory consolidation, in which local reactivations within the hippocampus during sharp-wave ripples are temporally aligned with neocortical excitability windows to enable transfer and integration of labile traces into distributed cortical networks [1,10,37].

The active systems consolidation model of memory propose that encoding occurs during waking, particularly through the hippocampus, whereas subsequent offline reactivations during nREM sleep promote stabilization and reorganization of memory traces in neocortex [38–40]. Here, we demonstrate that specific regions of this system (CA1d and CA1v) contribute differentially to this process. CA1d ripple–SO coupling in PFC and RSC was selectively associated with better performance in a spatial memory task, suggesting that dorsal hippocampal output during ripples are consistent with cortico-cortical plasticity for spatial representation. This observation aligns with anatomical and functional distinctions along the hippocampal longitudinal axis, with dorsal CA1 preferentially interacting with cortical networks involved in allocentric spatial representation and egocentric–allocentric transformations [20,41,42].

The selectivity of ripple–SO coupling effects to CA1d and not CA1v supports a functional dissociation in the contribution of hippocampal subregions to systems-level consolidation. While CA1v is interconnected with networks involved in affective processing, including PFC and ventral striatal areas [19,43,44], our data suggest that its output during ripples does not significantly modulate the neocortical dynamics critical for spatial discrimination in the current paradigm. This functional gradient is further reflected in the spectral and phase-locking patterns observed across regions, reinforcing the idea of topographically and temporally structured information transfer.

Our results resonate with human studies highlighting the role of sleep oscillations in memory consolidation. Indeed, previous work has established the behavioral relevance of SO–spindle–ripple triads in declarative memory retention, where temporal nesting between cortical SOs, thalamic spindles, and hippocampal ripples predicts memory performance [10,11,45]. Our rodent data corroborate and extend this framework by showing that not all ripple–spindle or ripple–SO events are equally effective. Those originating from CA1d and temporally coordinated with RSC or PFC rhythms are preferentially associated with successful memory retention. Furthermore, by subtracting baseline connectivity from pre-task sleep epochs, we isolate task-related network changes and demonstrate their predictive relationship with behavior, thus paralleling human EEG studies where sleep oscillatory reorganization is sensitive to memory demands [26,46].

In addition to SO and spindle coupling, we also find that CA1d–LEC ripple–spindle coupling relates to performance, suggesting that LEC, typically implicated in object-context encoding and spatial novelty [47,48], may also participate in memory trace integration during sleep. This is particularly relevant in the context of recent theories emphasizing the entorhinal cortex bidirectional role in modulating hippocampal activity and routing cortical inputs [49,50]. The temporal precision of ripple–spindle events between CA1d and LEC indicates that entorhinal processing is not merely feedforward or passive but may participate in reinstating and modifying representations during sleep.

Importantly, the ripple–SO and ripple–spindle effects reported here are temporally specific. Enhancements are constrained to defined windows relative to ripple onset and disappear outside those intervals. This suggests that hippocampo–cortical dialogue is not continuous but gated by precise oscillatory phase relationships. Such phase specificity is essential for spike-timing dependent plasticity and supports models in which cross-regional phase alignment enables synaptic consolidation [51–53].

Nevertheless, several limitations should be acknowledged in our study. First, the sample size, while appropriate for large-scale LFP, limits the granularity of behavioral correlations, especially when stratifying sessions by performance. Although median splits are standard in such contexts, future work using continuous regressors or larger datasets will offer greater sensitivity and resolution. Second, LFP-based measures provide coarse indices of network activity and cannot resolve fine-grained synaptic or circuit-level mechanisms. Our crosscorrelation analysis is a valuable tool, but inherently indirect. Third, although we report correlations between sleep connectivity and behavior, we do not establish causality. Disruption experiments targeting specific ripple–spindle motifs, perhaps via closed-loop optogenetics, would be essential to confirm functional relevance. Fourth, the spatial resolution of LFPs does not distinguish between local versus volume-conducted signals, and future studies will benefit from current source density or laminar recordings to address this.

Future research should pursue ensemble-level decoding of hippocampal replay events and assess their relationship to sleep oscillations and memory retention. Integrating calcium imaging or high-density unit recordings could help map the transformation of cell assemblies across sleep. Moreover, it will be critical to probe how neuromodulatory systems influence the temporal coordination of ripples, spindles, and SOs. Noradrenergic tone, for instance, is known to fluctuate across sleep stages and may modulate plasticity thresholds [54–56]. Finally, integrating this framework with disease models, such as Alzheimer’s disease, in which ripple–spindle coupling is impaired [57,58], could provide translational insights into cognitive decline and therapeutic interventions.

In summary, our study delineates a region- and oscillation-specific architecture for hippocampo–cortical coordination during sleep and their association with spatial memory consolidation. By combining behavioral assessment with temporally precise neural analyses, we identify ripple–spindle–SO motifs whose presence and strength predict learning outcomes, and we differentiate the contributions of CA1d and CA1v to this process. These findings refine existing models of memory systems consolidation and highlight how structured temporal coordination across brain regions underpins adaptive behavior.

## MATERIALS AND METHODS

### Animals

Ten adult male Sprague–Dawley rats (postnatal days 40–60; 300–350 g) were obtained from the Center for Innovation on Biomedical Experimental Models (CIBEM, Pontificia Universidad Católica de Chile). Animals were housed under controlled temperature (22 ± 1 °C) and a 12 h light/dark cycle (lights on at 08:00), with food and water available ad libitum. All experimental protocols were reviewed and approved by the Scientific Ethical Committee for the Care of Animals and the Environment of the Pontificia Universidad Católica (CEC-CAA), under approval number, protocol 220512003. All methods were carried out in accordance with relevant guidelines and regulations. This study is reported in compliance with the ARRIVE (Animal Research: Reporting of In Vivo Experiments) guidelines. Two weeks before surgery, animal handling was performed for 10 minutes daily for 3-5 consecutive days. Then, animals were habituated to the experimental arena and retention platform. For the arena, habituation was performed in the absence of objects. Animals were individually placed inside the arena facing the wall (alternating the entrance wall) and allowed to explore for 10 minutes for 3 consecutive days. For the retention platform habituation, animals spend 2-3 hours daily for 3 consecutive days, after exploring the arena.

### Object-place recognition task

The behavioral task was conducted once per day during the first six hours of the light phase (08:00–14:00), a period associated with a higher probability of nREM sleep [3]. We used aslightly modified version of the standard OPR task [3,31]. First, animals were placed to a retention platform for 60 min and allowed to sleep ad libitum (baseline sleep). Then, animals were transferred to the task arena. The arena consisted of a circular enclosure (65 cm diameter, 35 cm height) made of gray-painted PVC. Two identical objects were placed 15 cm from the arena walls. Object pairs were novel in every session and were used only once across the entire protocol, ensuring that animals were exposed to each object pair a single time. Object positions relative to distal cues, as well as the animal’s starting position, were randomized across phases and sessions. Rats were allowed to freely explore the arena for 10 min during the sample phase. Following exploration, animals were transferred back to the retention platform for 60 min and allowed to sleep ad libitum (task sleep). During the subsequent test phase, one object was displaced to the opposite quadrant of the arena, and the animal was returned to the center of the arena to explore. Animal behavior was video recorded throughout all sessions, and locomotor trajectories were extracted using DeepLabCut [59]. Each rat performed the task once per day, within a minimum of three consecutive experimental sessions per animal.

### Task scoring and discrimination index

Scoring was performed by quantifying the exploration time of each object using custom-developed software in MATLAB. Behavior was considered as exploration when the rat sniffed directly at the object from maximum distance of 2 cm. The exploration index was calculated using the exploration time of each object according to the equation:

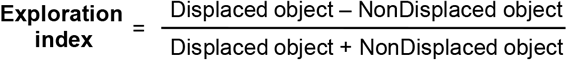

Positive values of the exploration index indicate preference to explore the displaced object (novelty condition) and suggest spatial memory formation, whereas negative values indicate preference to explore the non-displaced object (familiar condition) [30,31].

### Stereotaxic surgery for electrode implantation

Rats were anesthetized with isoflurane (4% for induction, 1.5–2% for maintenance) and positioned in a stereotaxic apparatus. Core body temperature was continuously monitored via a rectal probe and maintained at 36–37 °C throughout the surgical procedure (3–6 h) using a homeothermic blanket. After exposing the skull through a midline scalp incision, four craniotomies (∼1 mm diameter each) were performed in the right hemisphere at predetermined stereotaxic coordinates (CA1d: -4.0 AP, 2.0 ML, 2.7 DV; CA1v: -5.7 AP, -5.5 ML, 7.2 DV; PFC: 2.5 AP, -0.7 ML, 4.2 DV; RSC: -4.0 AP, 0.7 ML; LEC: −6.8 AP, 6.3 ML, 7.4 DV, 2.2 DV; all in mm). The dura mater was carefully removed to expose the cortical surface, which was kept moist with mineral oil to prevent tissue dehydration. Two additional craniotomies located anterior to bregma were made to accommodate the ground and reference electrodes. To further stabilize the implant, four supplementary craniotomies were prepared for anchoring skull screws, including two in the contralateral parietal bone, one in the ipsilateral parietal bone, and one positioned posterior to lambda. Following placement of the electrodes, craniotomies were sealed with silicone elastomer or bone wax, and the recording assembly was secured to the skull using dental acrylic. Postoperative care consisted of daily subcutaneous administration of enrofloxacin (5 mg/kg) and meloxicam (1 mg/kg) for three days. Animals were allowed to recover for a minimum of seven days prior to the onset of electrophysiological recordings.

### Electrophysiological recordings

Recordings were conducted during the light phase over four consecutive days in a Faraday-shielded enclosure. Animals were continuously recorded across the entire OPR task. Signals were acquired using Intan Technologies hardware (RHD system) at 20 kHz and synchronized with video recordings for behavioral monitoring. Data were converted to MATLAB format for offline analysis.

### Histology

After recordings, animals were anesthetized with isoflurane, electrolytically lesioned in each tetrode (10 μA positive current for 10 s was applied to every pair of channels), and allowed to recover for 48 hr. Then, rats were terminally anesthetized and intracardially perfused with a saline solution followed by a 20 min fixation with 4% paraformaldehyde. Brains were extracted and postfixed in paraformaldehyde overnight before being transferred to PBS-azide and sectioned coronally (60-80 μm slice thickness). Sections were further stained for Nissl substance. Locations of recording sites were performed under a light transmission microscope.

### Brain state classification

In addition to immobility, sleep scoring was performed using electrodes positioned in CA1d (to confirm theta oscillations [3]) and PFC (to confirm slow oscillations [60]). The LFP signals from these electrodes were downsampled to 1000 Hz, and time-frequency decomposition was performed with Fourier analysis using the LAN toolbox to obtain the signal’s power spectral density. The signal was analyzed for subsequent 10-second windows, where the raw LFP signal, the power spectrum, and the video recording were used to determine the stage of the sleep-wake cycle. One of three different stages was assigned to each window depending on the following criteria: if the power spectrum had a peak in slow wave activity (0.5-4 Hz) and the animal was immobile, the window was identified as nREM; if the power spectrum presented a peak in theta oscillations (4-10 Hz) and the animal was immobile, the window was identified as REM sleep; finally, if the animal was actively moving, the window was identified as awake. To assign a window into any of these categories, the criteria had to be fulfilled in at least 50% of the window. Exceptionally, a window was scored as undetermined if the previously described criteria were not met. This analysis was performed with a MATLAB (MathWorks) script.

### Detection of sleep oscillations

Theta oscillations were detected by calculating the continuous ratio between the envelope of theta (4–8 Hz) and delta waves (1–4 Hz) frequency bands filtered from the hippocampus LFP and computed by the Hilbert transform. A ratio of 1.4 standard deviation (SD) or higher during at least 2 s defined epochs of theta oscillations. For the detection of ripples, the LFP signal was downsampled to 1 kHz and band-pass filtered (100-250 Hz) using a zero-phase shift non-causal finite impulse filter with 0.5 Hz roll-off. Next, the signal was rectified, and low‐pass filtered at 20 Hz with a fourth-order Butterworth filter. This procedure yields a smooth envelope of the filtered signal, then a z-score normalized using the whole signal’s mean and SD. Epochs during which the normalized signal exceeded a 3 SD threshold and 50 ms of duration were considered ripples. The first point before the threshold that reached 1 SD was considered the ripple onset, and the first point after the threshold to reach 1 SD was considered the ripple end. The difference between onset and end of ripples was used to estimate the ripple duration. We introduced a 50 ms-refractory window to prevent double detections. To precisely determine the mean frequency, amplitude, and duration of each ripple, a spectral analysis using Morlet complex wavelets of seven cycles was performed. The protocol was adapted from a previously described method [17], and the LAN toolbox (http://lantoolbox.wikispaces.com) was used for the implementation. Spindles were detected in the downsampled hippocampal LFP signal (1 kHz), by calculating the maximum normalized wavelet power from the filtered signal (11-17 Hz). This signal was then Z-score normalized, using the whole signal’s mean and SD. Epochs during which the normalized signal exceeded a 1.4 SD threshold and 350 ms of duration were considered spindles. Spindles considered for analysis were exclusively detected during nREM epochs. Slow wave activity was detected by calculating the envelope of the slow oscillations (0.5–4 Hz) filtered from the PFC LFP and computed by the Hilbert transform. Slow wave activity was Z-scored during all sleep periods and epochs with amplitude 1.4 SD or higher, and at least 2 seconds were defined as slow wave activity.

### Peri-event cross-correlograms

Oscillatory events were cross-correlated by applying the ‘sliding-sweeps’ algorithm [61]. A time window of ± 1 s was defined with the 0-point assigned to the onset of a ripple. The timestamps of oscillatory events (ripples, spindles and Slow oscillations) within the time window were considered as a template and were represented by a vector relative to t = 0 s, with a time bin of 25 ms and normalized to the total number of events. Next, the window was shifted to successive events throughout the recording session, and an array of recurrences of templates was obtained.

### Statistical analysis

All statistical analyses were performed in MATLAB (MathWorks). Behavioral performance in the object–place recognition task was quantified using the discrimination index [3]. Associations between sleep architecture and behavior were assessed using Pearson or Spearman correlations as appropriate, and the relationship between nREM sleep duration and performance was further evaluated using linear mixed-effects models with nREM as a fixed effect and animal identity as a random intercept. Cross-regional coordination of sleep oscillations was quantified using peri-event cross-correlograms, aligning cortical spindle and slow oscillation events to the onset of hippocampal sharp-wave ripples detected in CA1d or CA1v. Event probability was computed in 25-ms bins within a ±1 s window relative to ripple onset, and cross-correlogram amplitude and latency were extracted within predefined temporal windows. Regional differences in coupling metrics were assessed using linear mixed-effects models with target region as a fixed effect and animal as a random intercept, with separate models for dorsal and ventral ripple sources. To assess performance-related modulation, task-induced changes in coupling were computed by subtracting baseline from task-related sleep measures and compared between low- and top-performing sessions (median split) using linear mixed-effects models. Non-parametric Wilcoxon tests were used for comparisons against zero and between groups when normality assumptions were violated, with false discovery rate correction applied where appropriate. Statistical significance was set at a two-tailed α = 0.05.

## Data availability

The datasets generated and analyzed during this study are available from the corresponding author upon reasonable request.

## Acknowledgments

This work was supported by FONDECYT grant 1230589 and Anillos ACT 210053. PF designed the study and wrote the manuscript; NE and MC performed experiments and analyses; GL, GF, AA, and AL-V contributed to data acquisition. The authors declare no competing interests. This manuscript was prepared with the assistance of AI-based tools (ChatGPT and Gemini). ChatGPT was used for improving the clarity, grammar, and overall flow of the text. Both AI-based tools were used for providing guidance on statistical analyses, including verification and refinement of the statistical treatment of results. In all cases, the authors thoroughly reviewed, edited, and validated any AI-generated content. AI-based tools were not used for core research tasks such as experimental design, data acquisition, primary data analysis, interpretation of results, or drawing scientific conclusions. The authors take full responsibility for all content of the manuscript.

