## Supplemental information for "Spatial memory performance is associated with region-specific coordination of hippocampo–cortical sleep oscillations"

### Supplementary Figures

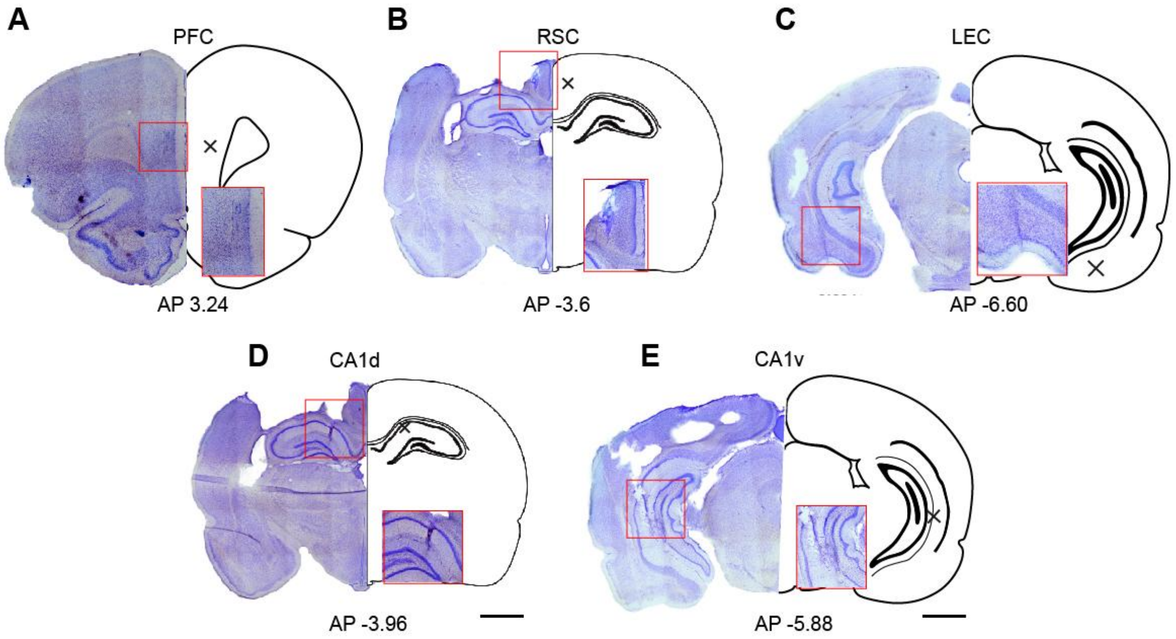

**Figure S1. Histological verification of electrode placement.** Representative Nissl-stained coronal brain sections illustrating the locations of recording electrodes in the prefrontal cortex (PFC; **A**), retrosplenial cortex (RSC; **B**), lateral entorhinal cortex (LEC; **C**), dorsal CA1 of the hippocampus (CA1d; **D**), and ventral CA1 of the hippocampus (CA1v; **E**). For each panel, the left image shows the histological section, and the right schematic depicts the corresponding atlas level. Red rectangles indicate the regions of interest and the approximate locations of electrode tracks or recording sites. Black crosses mark the estimated center of the recording area within each target structure. Anteroposterior (AP) coordinates relative to bregma are indicated below each panel. Scalebars, 2 mm.

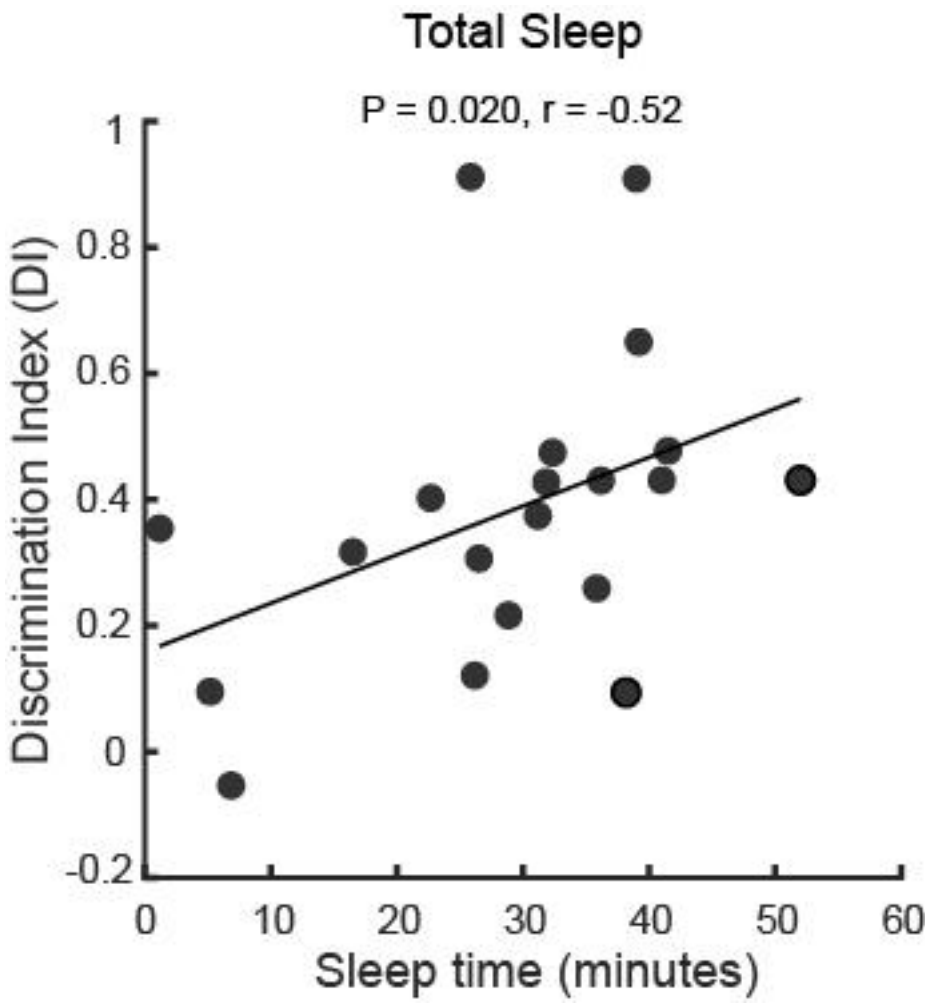

**Figure S2. Relationship between total sleep duration and spatial memory performance.** Scatter plot showing the association between total sleep time during the post-sample retention period and spatial memory performance, quantified by the discrimination index (DI) in the object–place recognition task. Each data point represents a single experimental session. The solid line indicates the linear regression fit, illustrating a positive relationship whereby longer total sleep duration is associated with higher discrimination indices, consistent with improved spatial memory performance (r= 0.52, P = 0.02, n = 10 rats, 20 sessions).

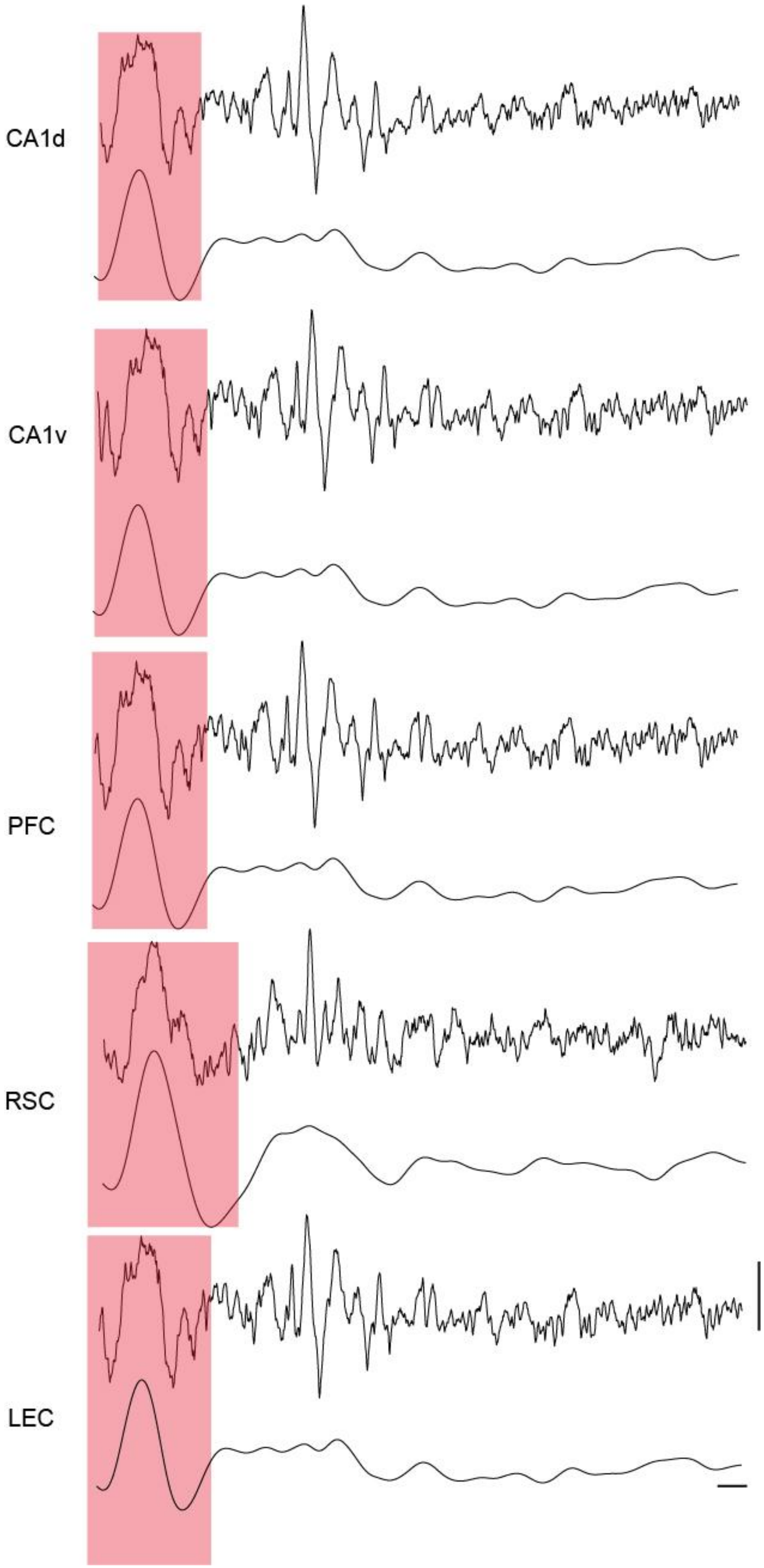
CA1d
CA1v
PFC
RSC
LEC

**Figure S3. Representative example of hippocampal–cortical slow oscillations during nREM sleep.** Example local field potential (LFP) traces recorded simultaneously from dorsal CA1 (CA1d), ventral CA1 (CA1v), prefrontal cortex (PFC), retrosplenial cortex (RSC), and lateral entorhinal cortex (LEC) during non–rapid eye movement (nREM) sleep. Red shaded regions indicate cortical slow oscillation (SO) events, illustrating the temporal alignment of large-amplitude low-frequency activity across hippocampal and cortical regions. For each channel, the raw LFP trace is shown above, with the corresponding low-frequency (0.1–4 Hz) SO component displayed below. This example highlights the widespread yet region-specific expression of slow oscillations across the hippocampo–cortical network during nREM sleep. Scale bars indicate 100 ms (horizontal) and 500 μV (vertical).

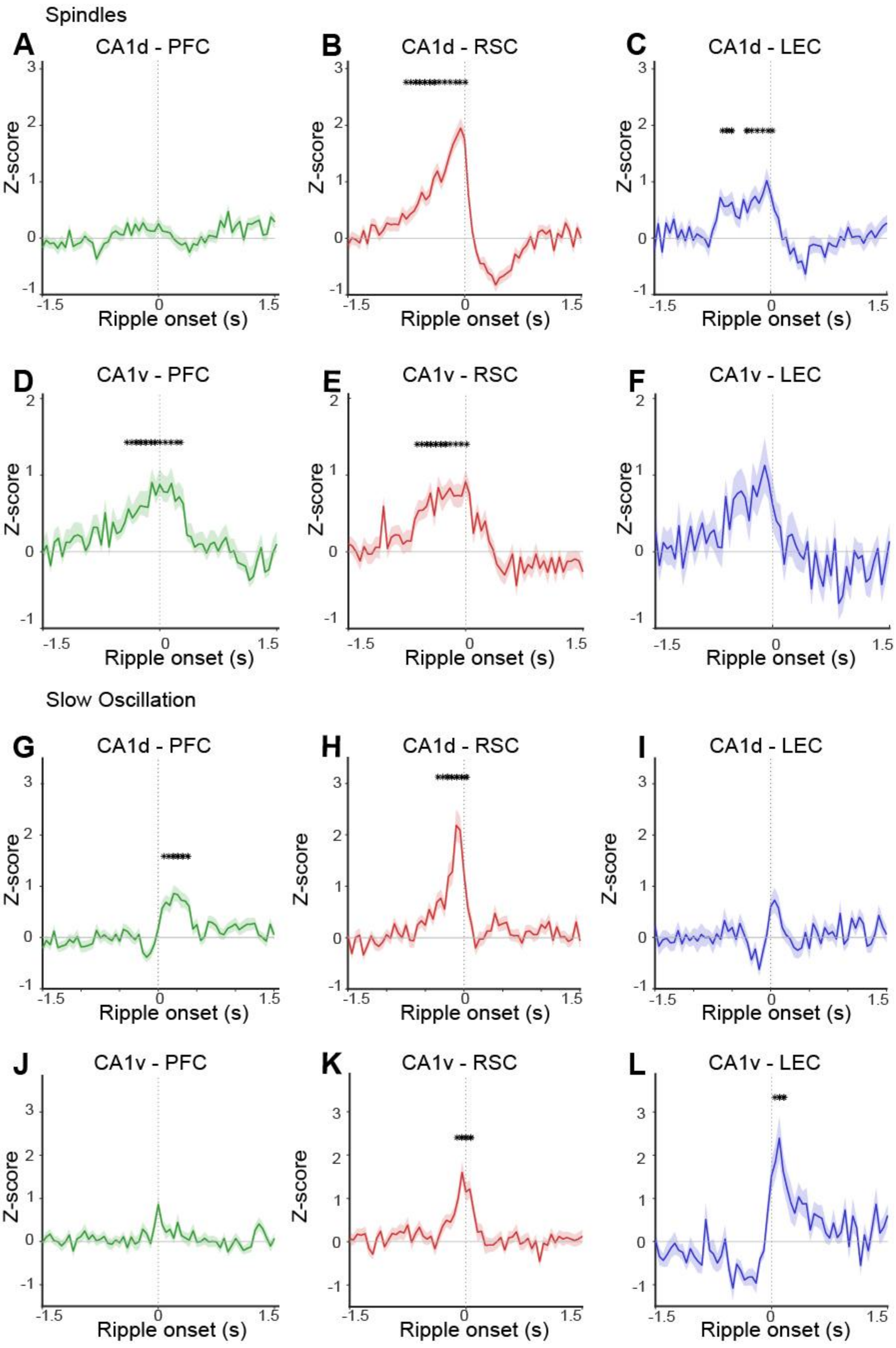
Spindles
A
CA1d - PFC
B
CA1d - RSC
C
CA1d - LEC
D
CA1v - PFC
E
CA1v - RSC
F
CA1v - LEC
Slow Oscillation
G
CA1d - PFC
H
CA1d - RSC
I
CA1d - LEC
J
CA1v - PFC
K
CA1v - RSC
L
CA1v - LEC
Z-score
Ripple onset (s)
-1.5
0
1.5

**Figure S4. Ripple-aligned cortical spindle and slow-oscillation probability across hippocampal–cortical pathways.** Average z-scored probability of cortical spindle (**A–F**) and slow-oscillation (SO; **G–L**) events aligned to hippocampal ripple onset (time 0) detected in dorsal CA1 (CA1d; **A–C, G–I**) or ventral CA1 (CA1v; **D–F, J–L**). Traces show mean ± s.e.m. across sessions for each hippocampal–cortical pair: prefrontal cortex (PFC, green), retrosplenial cortex (RSC, red), and lateral entorhinal cortex (LEC, blue). Vertical dashed lines indicate ripple onset. Asterisks denote time windows in which spindle or SO probability differed significantly from baseline (cluster-based or windowed statistical tests, $P < 0.05$). The panels illustrate pronounced region- and pole-specific modulation of cortical oscillations around hippocampal ripples, with strong pre- and peri-ripple increases in RSC and LEC relative to PFC, and distinct profiles for CA1d- versus CA1v-aligned events.

### Supplementary Tables

| | | Sample phase exploration time (s) | | | Test phase exploration time (s) | | |
|---|---|---|---|---|---|---|---|
| **Animal** | **Session** | **obj 1 (fixed)** | **obj2 (fixed)** | **total time** | **obj1 (fixed)** | **obj2 (displaced )** | **total time** |
| AA04 | Day02 | 21.6 | 29.2 | 50.7 | 18.7 | 22.5 | 41.2 |
| AA04 | Day04 | 36.3 | 41.7 | 78.0 | 9.3 | 50.3 | 59.6 |
| AA06 | Day01 | 26.6 | 28.4 | 55.0 | 17.9 | 27.7 | 45.6 |
| AA06 | Day04 | 53.0 | 38.3 | 91.3 | 19.9 | 29.2 | 49.1 |
| AA07 | Day01 | 65.6 | 40.9 | 106.6 | 47.2 | 43.9 | 91.1 |
| AA07 | Day02 | 71.9 | 88.7 | 160.6 | 40.7 | 55.7 | 96.4 |
| AA07 | Day04 | 62.9 | 48.4 | 111.4 | 24.8 | 58.8 | 83.6 |
| AA12 | Day01 | 45.9 | 34.8 | 80.7 | 25.7 | 44.3 | 70.0 |
| AA12 | Day04 | 31.2 | 32.7 | 63.9 | 15.3 | 38.5 | 53.9 |
| PF24 | Day01 | 36.7 | 48.3 | 85.0 | 23.9 | 60.0 | 83.9 |
| PF24 | Day02 | 38.9 | 30.4 | 69.2 | 13.3 | 35.1 | 48.4 |
| PF26 | Day01 | 63.5 | 39.8 | 103.3 | 27.1 | 34.9 | 62.0 |
| PF26 | Day02 | 72.7 | 91.9 | 164.6 | 44.1 | 80.3 | 124.4 |
| PF28 | Day01 | 27.0 | 15.7 | 42.7 | 38.3 | 61.0 | 99.3 |
| PF28 | Day02 | 49.1 | 64.7 | 113.8 | 15.3 | 29.3 | 44.6 |
| PF31 | Day04 | 25.8 | 20.6 | 46.5 | 11.3 | 28.3 | 39.6 |
| PF33 | Day01 | 33.3 | 36.1 | 69.5 | 8.5 | 41.3 | 49.8 |
| PF33 | Day02 | 27.7 | 22.0 | 49.7 | 3.1 | 15.2 | 18.3 |
| PF44 | Day01 | 63.6 | 51.1 | 114.8 | 15.3 | 34.7 | 50.0 |
| PF44 | Day02 | 35.8 | 30.6 | 66.5 | 26.0 | 46.4 | 72.4 |
| **Mean** | | 44.5 | 41.7 | 86.2 | 21.6 | 41.9 | 63.5 |
| **SD** | | 17.0 | 20.1 | 34.9 | 13.1 | 15.7 | 26.4 |

**Table S1**. **Individual exploration times in the object-place recognition task**. Table presents the exploration times (in seconds) for each animal and session during the sample and test phases of the task. Columns detail the time spent exploring the two identical objects during the sample phase (obj 1, obj 2) and the total exploration time. For the test phase, exploration times are shown for the familiar condition (obj 1, fixed) and the novelty condition (obj 2, displaced), along with the total exploration time. Mean and standard deviation (SD) values across all sessions are provided at the bottom. The higher average exploration time for the displaced object (41.9 s) compared to the fixed object (21.6 s) indicates a group-level preference for the novel spatial configuration, reflecting successful spatial memory retrieval.

| Baseline sleep | | | | | | | |
|---|---|---|---|---|---|---|---|
| **Animal** | **Session** | **QW (min)** | **nREM (min)** | **REM (min)** | **QW (%)** | **nREM (%)** | **REM (%)** |
| AA04 | Day02 | 32.8 | 24.8 | 3.0 | 54.1 | 40.9 | 4.9 |
| AA04 | Day04 | 35.8 | 18.2 | 4.8 | 60.9 | 30.9 | 8.2 |
| AA06 | Day01 | 37.8 | 17.5 | 3.0 | 64.9 | 30.0 | 5.1 |
| AA06 | Day04 | 24.8 | 28.7 | 7.2 | 40.9 | 47.3 | 11.8 |
| AA07 | Day01 | 57.0 | 0.0 | 0.0 | 100.0 | 0.0 | 0.0 |
| AA07 | Day02 | 46.8 | 11.7 | 0.3 | 79.6 | 19.8 | 0.6 |
| AA07 | Day04 | 29.5 | 22.2 | 6.7 | 50.6 | 38.0 | 11.4 |
| AA12 | Day01 | 60.0 | 0.0 | 0.0 | 100.0 | 0.0 | 0.0 |
| AA12 | Day04 | 28.7 | 27.0 | 4.7 | 47.5 | 44.8 | 7.7 |
| PF24 | Day01 | 21.2 | 38.7 | 0.5 | 35.1 | 64.1 | 0.8 |
| PF24 | Day02 | 29.5 | 31.0 | 0.0 | 48.8 | 51.2 | 0.0 |
| PF26 | Day01 | 42.8 | 18.8 | 0.0 | 69.5 | 30.5 | 0.0 |
| PF26 | Day02 | 46.8 | 11.7 | 0.0 | 80.1 | 19.9 | 0.0 |
| PF28 | Day01 | 24.0 | 32.0 | 3.7 | 40.2 | 53.6 | 6.1 |
| PF28 | Day02 | 20.5 | 38.3 | 2.0 | 33.7 | 63.0 | 3.3 |
| PF31 | Day04 | 27.7 | 29.2 | 3.0 | 46.2 | 48.7 | 5.0 |
| PF33 | Day01 | 35.8 | 26.7 | 0.0 | 57.3 | 42.7 | 0.0 |
| PF33 | Day02 | 33.8 | 25.7 | 0.0 | 56.9 | 43.1 | 0.0 |
| PF44 | Day01 | 51.0 | 8.8 | 0.0 | 85.2 | 14.8 | 0.0 |
| PF44 | Day02 | 31.5 | 28.5 | 2.0 | 50.8 | 46.0 | 3.2 |
| **Mean** | | 35.9 | 22.0 | 2.0 | 60.1 | 36.5 | 3.4 |
| **SD** | | 11.5 | 11.1 | 2.4 | 19.9 | 18.2 | 4.0 |

| Task sleep | | | | | | | |
|---|---|---|---|---|---|---|---|
| **Animal** | **Session** | **QW (min)** | **nREM (min)** | **REM (min)** | **QW (%)** | **nREM (%)** | **REM (%)** |
| AA04 | Day02 | 21.8 | 29.7 | 8.5 | 36.4 | 49.4 | 14.2 |
| AA04 | Day04 | 20.3 | 32.7 | 6.5 | 34.2 | 54.9 | 10.9 |
| AA06 | Day01 | 28.2 | 26.7 | 2.2 | 49.4 | 46.8 | 3.8 |
| AA06 | Day04 | 31.7 | 22.3 | 3.8 | 54.8 | 38.6 | 6.6 |
| AA07 | Day01 | 50.0 | 6.8 | 0.0 | 88.0 | 12.0 | 0.0 |
| AA07 | Day02 | 53.0 | 5.2 | 0.0 | 91.1 | 8.9 | 0.0 |
| AA07 | Day04 | 46.5 | 22.3 | 0.3 | 67.2 | 32.3 | 0.5 |
| AA12 | Day01 | 53.8 | 1.2 | 0.0 | 97.9 | 2.1 | 0.0 |
| AA12 | Day04 | 6.8 | 41.5 | 10.5 | 11.6 | 70.5 | 17.8 |
| PF24 | Day01 | 19.3 | 40.3 | 0.7 | 32.0 | 66.9 | 1.1 |
| PF24 | Day02 | 25.3 | 36.2 | 0.0 | 41.2 | 58.8 | 0.0 |
| PF26 | Day01 | 43.2 | 16.2 | 0.3 | 72.3 | 27.1 | 0.6 |
| PF26 | Day02 | 34.2 | 26.0 | 0.5 | 56.3 | 42.9 | 0.8 |

| | | | | | | | |
|---|---|---|---|---|---|---|---|
| PF28 | Day01 | 29.0 | 27.7 | 3.5 | 48.2 | 46.0 | 5.8 |
| PF28 | Day02 | 34.3 | 22.3 | 3.5 | 57.1 | 37.1 | 5.8 |
| PF31 | Day04 | 27.8 | 26.5 | 5.3 | 46.6 | 44.4 | 8.9 |
| PF33 | Day01 | 27.5 | 32.3 | 0.0 | 46.0 | 54.0 | 0.0 |
| PF33 | Day02 | 20.8 | 36.8 | 2.2 | 34.8 | 61.6 | 3.6 |
| PF44 | Day01 | 18.0 | 36.3 | 5.2 | 30.3 | 61.1 | 8.7 |
| PF44 | Day02 | 27.3 | 30.8 | 5.0 | 43.3 | 48.8 | 7.9 |
| **Mean** | | 31.0 | 26.0 | 2.9 | 51.9 | 43.2 | 4.9 |
| **SD** | | 12.7 | 11.4 | 3.1 | 22.1 | 19.0 | 5.3 |

**Table S2**. **Sleep architecture during baseline and task-related resting periods**. Table shows the time (in minutes) and percentage of total time spent in Quiet Wakefulness (QW), non-REM (nREM) sleep, and Rapid Eye Movement (REM) sleep for each animal and session while animals were in the retention platform. Data are stratified by recording epoch: Baseline sleep and task sleep. Mean and standard deviation (SD) values across all sessions are provided at the bottom. The data confirms that animals engaged in substantial sleep during the retention periods.

| Animal | CA1d | CA1v | PFC | RSC | LEC |
|---|---|---|---|---|---|
| AA04 | + | - | + | + | + |
| AA06 | + | + | + | + | + |
| AA07 | + | + | + | + | + |
| AA12 | + | + | + | + | - |
| PF24 | + | + | + | + | - |
| PF26 | + | + | + | + | - |
| PF28 | + | - | + | - | + |
| PF31 | + | + | + | + | - |
| PF33 | + | + | + | + | + |
| PF39 | + | + | + | + | - |
| PF40 | + | + | + | + | - |
| PF44 | + | - | + | + | + |

**Table S3**. **Summary of electrode locations across recorded brain areas**. Table shows the availability of viable local field potential recordings for each animal across the targeted brain areas: dorsal and ventral hippocampus (CA1d, CA1v), prefrontal cortex (PFC), retrosplenial cortex (RSC), and lateral entorhinal cortex (LEC). A plus sign (+) represents histologically confirmed electrode location and inclusion of the signals in the data analysis, while a minus sign (-) indicates that data from that specific area was unavailable due to missed targeting.

**Discrimination Index ~ nREM + REM + (1|animal)**

| Predictor | Estimate | SE | t | df | P value |
|---|---|---|---|---|---|
| (Intercept) | 0.1222 | 0.1203 | 1.0160 | 17 | 0.3239 |
| nREM | 0.0112 | 0.0048 | 2.3200 | 17 | 0.0330 |
| REM | -0.0114 | 0.0176 | -0.6494 | 17 | 0.5248 |

| CI lower | CI upper | AIC | BIC | Log likelihood | residual variance |
|---|---|---|---|---|---|
| -0.1316 | 0.3760 | 4.3647 | 9.3433 | 2.8177 | 0.0442 |
| 0.0010 | 0.0215 | 4.3647 | 9.3433 | 2.8177 | 0.0442 |
| -0.0486 | 0.0257 | 4.3647 | 9.3433 | 2.8177 | 0.0442 |

**Table S4. Linear mixed-effects model relating spatial memory performance to nREM and REM sleep duration.** The table summarizes results from a multivariable linear mixed-effects model assessing the association between non–rapid eye movement (nREM) and rapid eye movement (REM) sleep duration with spatial memory performance, quantified by the discrimination index. The model included both nREM and REM sleep duration as fixed effects and a random intercept for each Animal to account for repeated measurements. Fixed-effect estimates for the Intercept, nREM, and REM sleep duration are reported together with their standard errors (SE), t-statistics (t), degrees of freedom (df), p-values, and 95% confidence intervals (CI). Measures of model fit, including Akaike information criterion (AIC), Bayesian information criterion (BIC), log-likelihood, and residual variance, are also provided. The analysis confirms a significant positive association for nREM sleep ($P = 0.033$) while REM sleep shows no significant effect ($P = 0.52$).

**Crosscorrelagram amplitude ~ target area + (1|animal)**

**Crosscorrelagram latency ~ target area + (1|animal)**

| Source | Metric | Predictor | Estimate | SE | t | df | P value |
|---|---|---|---|---|---|---|---|
| CA1d | Amplitude | (Intercept) | 2.55 | 0.25 | 10.38 | 125 | 0.000000 |
| CA1d | Amplitude | target-RSC | 0.92 | 0.18 | 5.23 | 125 | 0.000001 |
| CA1d | Latency | (Intercept) | -289.99 | 48.48 | -5.98 | 125 | 0.000000 |
| CA1d | Latency | target-RSC | 24.38 | 47.03 | 0.52 | 125 | 0.605001 |
| CA1v | Amplitude | (Intercept) | 2.67 | 0.23 | 11.60 | 132 | 0.000000 |
| CA1v | Amplitude | target-RSC | -0.14 | 0.10 | -1.33 | 132 | 0.187415 |
| CA1v | Latency | (Intercept) | -80.60 | 26.88 | -3.00 | 132 | 0.003243 |
| CA1v | Latency | target-RSC | -193.28 | 38.01 | -5.08 | 132 | 0.000001 |

| CI lower | CI upper | AIC | BIC | Log likelihood | Residual variance |
|---|---|---|---|---|---|
| 2.06 | 3.04 | 334.67 | 346.04 | -163.33 | 0.63 |
| 0.57 | 1.26 | 334.67 | 346.04 | -163.33 | 0.63 |
| -385.93 | -194.05 | 1756.18 | 1767.56 | -874.09 | 50102.19 |
| -68.69 | 117.46 | 1756.18 | 1767.56 | -874.09 | 50102.19 |
| 2.21 | 3.12 | 277.62 | 289.21 | -134.81 | 0.36 |
| -0.34 | 0.07 | 277.62 | 289.21 | -134.81 | 0.36 |
| -133.77 | -27.43 | 1833.79 | 1845.38 | -912.89 | 48407.22 |
| -268.48 | -118.09 | 1833.79 | 1845.38 | -912.89 | 48407.22 |

**Table S5**. **Linear Mixed-Effects Models of cortical spindle modulation by hippocampal ripples.** This table summarizes two separate linear mixed-effects models analyzing the amplitude and latency of ripple-triggered cortical spindles. The models predict these metrics based on the cortical target region, with source indicating the hippocampal origin of the ripples (CA1d or CA1v). A random intercept for each animal is included. Estimates for the intercept (reference region) and the difference for the comparison region (Target_RSC) are shown with standard errors (SE), t-statistics, degrees of freedom (df), p-values, and 95% confidence intervals (CI). Model fit statistics are provided.

**Crosscorrelagram amplitude ~ target area + (1|animal)**

**Crosscorrelagram latency ~ target area + (1|animal)**

| Source | Metric | Predictor | Estimate | SE | t | df | P value |
|---|---|---|---|---|---|---|---|
| CA1d | Amplitude | (Intercept) | 2.34 | 0.31 | 7.55 | 146 | 0.000000 |
| CA1d | Amplitude | Target-RSC | 0.98 | 0.24 | 4.13 | 146 | 0.000061 |
| CA1d | Latency | (Intercept) | 199.40 | 12.61 | 15.82 | 146 | 0.000000 |
| CA1d | Latency | Target-RSC | -334.01 | 19.02 | -17.56 | 146 | 0.000000 |
| CA1v | Amplitude | (Intercept) | 4.19 | 0.49 | 8.50 | 64 | 0.000000 |
| CA1v | Amplitude | Target-RSC | -1.47 | 0.41 | -3.60 | 64 | 0.000625 |
| CA1v | Latency | (Intercept) | 82.35 | 17.50 | 4.70 | 64 | 0.000014 |
| CA1v | Latency | Target-RSC | -101.74 | 20.32 | -5.01 | 64 | 0.000005 |

| CI lower | CI upper | AIC | BIC | Log likelihood | Residual variance |
|---|---|---|---|---|---|
| 1.73 | 2.95 | 547.91 | 559.90 | -269.95 | 1.94 |
| 0.51 | 1.46 | 547.91 | 559.90 | -269.95 | 1.94 |
| 174.48 | 224.31 | 1832.11 | 1844.10 | -912.06 | 13189.77 |
| -371.61 | -296.42 | 1832.11 | 1844.10 | -912.06 | 13189.77 |
| 3.20 | 5.17 | 228.99 | 237.75 | -110.49 | 1.32 |
| -2.29 | -0.66 | 228.99 | 237.75 | -110.49 | 1.32 |
| 47.38 | 117.32 | 760.14 | 768.89 | -376.07 | 5208.90 |
| -142.33 | -61.16 | 760.14 | 768.89 | -376.07 | 5208.90 |

**Table S6**. **Linear Mixed-Effects Models of cortical slow oscillation modulation by hippocampal ripples.** This table summarizes two separate linear mixed-effects models analyzing the amplitude and latency of ripple-triggered cortical slow oscillations. The models predict these metrics based on the cortical target region, with source indicating the hippocampal origin of the ripples (CA1d or CA1v). A random intercept for each animal is included. Estimates for the intercept (reference region) and the difference for the comparison region (Target_RSC) are shown with standard errors (SE), t-statistics, degrees of freedom (df), p-values, and 95% confidence intervals (CI). Model fit statistics are provided.

**Delta sleep (task – baseline) ~ group + (1|animal)**

| Pair | Window (ms) | Predictor | Estimate | SE | t | df | P value |
|---|---|---|---|---|---|---|---|
| CA1d-RSC | -800 to +50 | (Intercept) | 0.052 | 0.098 | 0.528 | 16 | 0.604 |
| CA1d-RSC | -800 to +50 | Group-Top | 0.022 | 0.131 | 0.163 | 16 | 0.872 |
| CA1d-LEC | -350 to 0 | (Intercept) | 0.091 | 0.1405 | 0.649 | 11 | 0.529 |
| CA1d-LEC | -350 to 0 | Group-Top | 0.485 | 0.2068 | 2.346 | 11 | 0.0387 |
| CA1v-RSC | -650 to +50 | (Intercept) | 0.007 | 0.127 | 0.058 | 12 | 0.954 |
| CA1v-RSC | -650 to +50 | Group-Top | 0.237 | 0.180 | 1.314 | 12 | 0.213 |
| CA1v-PFC | -450 to +300 | (Intercept) | -0.060 | 0.118 | -0.50 | 12 | 0.621 |
| CA1v-PFC | -450 to +300 | Group-Top | 0.127 | 0.132 | 0.959 | 12 | 0.356 |

| CI lower | CI upper | AIC | BIC | Log likelihood | Residual variance |
|---|---|---|---|---|---|
| -0.157 | 0.262 | 13.349 | 16.911 | -2.675 | 0.066 |
| -0.257 | 0.300 | 13.349 | 16.911 | -2.675 | 0.066 |
| -0.218 | 0.4005 | 19.164 | 21.424 | -5.582 | 0.1382 |
| 0.0302 | 0.9406 | 19.164 | 21.424 | -5.582 | 0.1382 |
| -0.271 | 0.285 | 17.320 | 19.876 | -4.660 | 0.114 |
| -0.156 | 0.630 | 17.320 | 19.876 | -4.660 | 0.114 |
| -0.320 | 0.199 | 8.730 | 11.286 | -0.365 | 0.031 |
| -0.161 | 0.414 | 8.730 | 11.286 | -0.365 | 0.031 |

**Table S7. Linear Mixed-Effects Models of task-related changes in hippocampo-cortical spindles coupling.** Results from linear mixed-effects models evaluating whether task performance (Group: Top vs. Low performers) predicts the change in sleep coupling strength (Delta sleep [task – baseline]). Separate models were run for specific pair connections (CA1d-RSC) and time windows relative to ripple onset. Each model includes Group as a fixed effect and a random intercept for Animal. The table lists estimates for the Intercept (baseline change) and the effect of being in the Top performance group, along with SE, t-statistics, df, p-values, and 95% CIs. Model fit statistics (AIC, BIC, Log Likelihood, Residual Variance) are included.

**Delta sleep (task – baseline) ~ group + (1|animal)**

| Pair | Window | Predictor | Estimate | SE | t | df | P value |
|---|---|---|---|---|---|---|---|
| CA1d-PFC | +50 to +400 | (Intercept) | -0.273 | 0.252 | -1.08 | 16 | 0.29 |
| CA1d-PFC | +50 to +400 | Group-Top | 0.577 | 0.240 | 2.40 | 16 | 0.028 |
| CA1d-RSC | -350 to +50 | (Intercept) | -0.212 | 0.216 | -0.98 | 14 | 0.34 |
| CA1d-RSC | -350 to +50 | Group-Top | 0.922 | 0.183 | 5.02 | 14 | 0.00019 |
| CA1v-LEC | 0 to +200 | (Intercept) | -0.137 | 0.271 | -0.50 | 4 | 0.63 |
| CA1v-LEC | 0 to +200 | Group-Top | -0.301 | 0.384 | -0.78 | 4 | 0.47 |
| CA1v-RSC | -150 to +100 | (Intercept) | 0.270 | 0.144 | 1.8 | 10 | 0.091 |
| CA1v-RSC | -150 to +100 | Group-Top | -0.405 | 0.204 | -1.9 | 10 | 0.07567 |

| CI lower | CI upper | AIC | BIC | Log likelihood | Residual variance |
|---|---|---|---|---|---|
| -0.807 | 0.261 | 39.296 | 42.858 | -15.648 | 0.109 |
| 0.069 | 1.085 | 39.296 | 42.858 | -15.648 | 0.109 |
| -0.674 | 0.251 | 25.826 | 28.917 | -8.913 | 0.041 |
| 0.528 | 1.315 | 25.826 | 28.917 | -8.913 | 0.041 |
| -0.891 | 0.616 | 15.965 | 15.132 | -3.982 | 0.221 |
| -1.366 | 0.765 | 15.965 | 15.132 | -3.982 | 0.221 |
| -0.052 | 0.592 | 17.117 | 19.057 | -4.559 | 0.125 |
| -0.860 | 0.050 | 17.117 | 19.057 | -4.559 | 0.125 |

**Table S8**. **Linear Mixed-Effects Models of task-related changes in hippocampo-cortical slow oscillations coupling.** Results from linear mixed-effects models evaluating whether task performance (Group: Top vs. Low performers) predicts the change in sleep coupling strength (Delta sleep [task – baseline]). Separate models were run for specific pair connections (CA1d-RSC) and time windows relative to ripple onset. Each model includes Group as a fixed effect and a random intercept for Animal. The table lists estimates for the Intercept (baseline change) and the effect of being in the Top performance group, along with SE, t-statistics, df, p-values, and 95% CIs. Model fit statistics (AIC, BIC, Log Likelihood, Residual Variance) are included.
